# Ref-1/APE1 inhibition with novel small molecules blocks ocular neovascularization

**DOI:** 10.1101/296590

**Authors:** Sardar Pasha Sheik Pran Babu, Kamakshi Sishtla, Rania S. Sulaiman, Bomina Park, Trupti Shetty, Fenil Shah, Melissa L. Fishel, James H. Wikel, Mark R. Kelley, Timothy W. Corson

## Abstract

Ocular neovascular diseases like wet age-related macular degeneration are a major cause of blindness. Novel therapies are greatly needed for these diseases. One appealing antiangiogenic target is reduction-oxidation factor 1-apurinic/apyrimidinic endonuclease 1 (Ref-1/APE1). This protein can act as a redox-sensitive transcriptional activator for NF-κB and other pro-angiogenic transcription factors. An existing inhibitor of Ref-1’s function, APX3330, previously showed antiangiogenic effects. Here, we developed improved APX3330 derivatives and assessed their antiangiogenic activity. We synthesized APX2009 and APX2014 and demonstrated enhanced inhibition of Ref-1 function in a DNA-binding assay compared to APX3330. Both compounds were antiproliferative against human retinal microvascular endothelial cells (HRECs; GI50 APX2009: 1.1 μM, APX2014: 110 nM) and macaque choroidal endothelial cells (Rf/6a GI_50_ APX2009: 26 μM, APX2014: 5.0 μM). Both compounds significantly reduced the ability of HRECs and Rf/6a cells to form tubes at mid nanomolar concentrations compared to control, and both significantly inhibited HREC and Rf/6a cell migration in a scratch wound assay, reducing NF-κB activation and downstream targets. *Ex vivo*, both APX2009 and APX2014 inhibited choroidal sprouting at low micromolar and high nanomolar concentrations respectively. In the laser-induced choroidal neovascularization mouse model, intraperitoneal APX2009 treatment significantly decreased lesion volume by 4-fold compared to vehicle (*p* < 0.0001, ANOVA with Dunnett’s post hoc tests), without obvious intraocular or systemic toxicity. Thus, Ref-1 inhibition with APX2009 and APX2014 blocks ocular angiogenesis *in vitro* and *ex vivo*, and APX2009 is an effective systemic therapy for CNV *in vivo*, establishing Ref-1 inhibition as a promising therapeutic approach for ocular neovascularization.

## Introduction

Ocular neovascularization is the key pathobiological feature of diseases like proliferative diabetic retinopathy (PDR), retinopathy of prematurity (ROP), and wet age-related macular degeneration (AMD), which together are major causes of blindness (Campochiaro, 2013). In PDR and ROP, abnormal blood vessels grow in and on the retina, while in wet AMD, neovessels grow from the pigmented, subretinal choroid layer into the retina. In all cases, neovessels disrupt retinal architecture and can hemorrhage, leading to blindness. Although the exact stimuli promoting neovascularization are not always well characterized, hypoxia and inflammation both play crucial roles. The currently used, FDA approved pharmacological treatments for these diseases are all biologics targeting the vascular endothelial growth factor (VEGF) signaling pathway, such as ranibizumab and aflibercept (Prasad et al., 2010). Although these therapeutic agents have been very successful, significant proportions of patients are resistant and refractory (Lux et al., 2007; Falavarjani and Nguyen, 2013). Moreover, serious side effects including hemorrhage and endophthalmitis are possible. Therefore, development of novel therapeutic approaches targeting other signaling pathways is crucial.

One such potential target is the reduction-oxidation factor 1-apurinic/apyrimidinic endonuclease 1 (Ref-1/APE1), an intracellular signaling nexus with important roles in transducing proangiogenic stimuli. This bifunctional protein has an endonuclease role essential for base excision repair (APE1), while the Ref-1 activity is a redox-sensitive transcriptional activator (Shah et al., 2017). Ref-1 redox signaling is a highly regulated process that reduces oxidized cysteine residues in specific transcription factors as part of their transactivation (Xanthoudakis and Curran, 1992; Xanthoudakis et al., 1992; Evans et al., 2000; Lando et al., 2000; Nishi et al., 2002; Seo et al., 2002; Li et al., 2010; Fishel et al., 2011; Cardoso et al., 2012; Kelley et al., 2012; Luo et al., 2012; Zhang et al., 2013; Fishel et al., 2015; Logsdon et al., 2016). This redox signaling affects numerous transcription factors including HIF-1α, NF-κB, and others. The regulation of HIF-1α and NF-κB are particularly relevant to angiogenesis and eye diseases (Evans et al., 2000; Nishi et al., 2002; Seo et al., 2002; Fishel et al., 2011; Cardoso et al., 2012; Fishel et al., 2015; Logsdon et al., 2016).

Excitingly, Ref-1 activity can be targeted pharmacologically. APX3330 (formerly called E3330) is a specific Ref-1/APE1 redox inhibitor. APX3330 has been extensively characterized as a direct, highly selective inhibitor of Ref-1 redox activity that does not affect the protein’s endonuclease activity (Luo et al., 2008; Fishel et al., 2010; Su et al., 2011; Cardoso et al., 2012; Luo et al., 2012; Zhang et al., 2013; Fishel et al., 2015). Pharmacologic inhibition of Ref-1 via APX3330 blocks Ref-1 redox activity on NF-κB, HIF-1α, AP-1, and STAT3, decreasing transcription factor binding to DNA *in vitro* (Fishel et al., 2011; Jedinak et al., 2011; Luo et al., 2012). APX3330 has entered Phase I clinical trials for safety and recommended Phase II dose (RP2D) in cancer patients (NCT03375086); however, the safety and dose administration of APX3330 have been previously established by Eisai Inc. through a prior development program for a non-cancer, Hepatitis C indication that evaluated the preclinical toxicology, Phase I and Phase II safety and clinical profile in more than 400 non-cancer patients (Shah et al., 2017).

Ref-1/APE1 is highly expressed during retinal development, and in retinal pigment epithelium (RPE) cells, pericytes, choroidal endothelial cells and retinal endothelial cells (Chiarini et al., 2000; Jiang et al., 2011; Li et al., 2014a). More generally, Ref-1 is frequently upregulated in regions of tissues in which inflammation is present (Zou et al., 2009; Kelley et al., 2010). APX3330 was previously shown to block *in vitro* angiogenesis, as evidenced by proliferation, migration, and tube formation of retinal and choroidal endothelial cells (Jiang et al., 2011; Li et al., 2014b). Indeed, APX3330 delivered intravitreally (directly into the eye) reduced neovascularization in the very low density lipoprotein receptor (VLDLR) knockout mouse model of retinal neovascularization (Jiang et al., 2011), and also in laser-induced choroidal neovascularization (L-CNV) (Li et al., 2014b), the most widely used animal model that recapitulates features of wet AMD (Grossniklaus et al., 2010).

While the lead clinical candidate is effective in preclinical cancer studies, we also sought novel, second generation Ref-1 inhibitors that would have increased efficacy in antiangiogenic and anti-inflammatory transcription factor (NF-κB, HIF-1α) inhibition, as well as new chemical properties. We present here the synthesis of APX3330 derivatives APX2009 (reported previously as a neuroprotective agent (Kelley et al., 2016)), and APX2014 (reported here for the first time (Kelley and Wikel, 2015)). We go on to show that these compounds have antiangiogenic activity against retinal and choroidal endothelial cells both in culture and in an *ex vivo* choroidal sprouting model. Finally, we show that APX2009 is an effective *systemic* treatment for L-CNV, establishing these compounds as leads for further development for neovascular eye diseases.

## Materials and Methods

### Synthetic Methods

The compounds were synthesized by Cascade Custom Chemistry (Eugene, OR) and provided by Apexian Pharmaceuticals. In summary (Fig. 1A), the common intermediate iodolawsone (2-iodo-3-hydroxy-1,4-naphthoquinone (**1**); available from Cascade Custom Chemistry) was treated with 2-propylacrylic acid (**2**), with oxalyl chloride and the corresponding amine, and with sodium methoxide in methanol to yield (*E*)-*N*,*N*-diethyl-2-((3-methoxy-1,4-dioxo-1,4-dihydronaphthalen-2-yl)methylene)pentanamide (**6a**; APX2009), and (*E*)-*N*-methoxy-2-((3-methoxy-1,4-dioxo-1,4-dihydronaphthalen-2-yl)methylene)pentanamide (**6b**; APX2014). Full synthetic details can be found in the Supplemental Methods. APX3330 was synthesized as described (Luo et al., 2008).

**Fig. 1.**
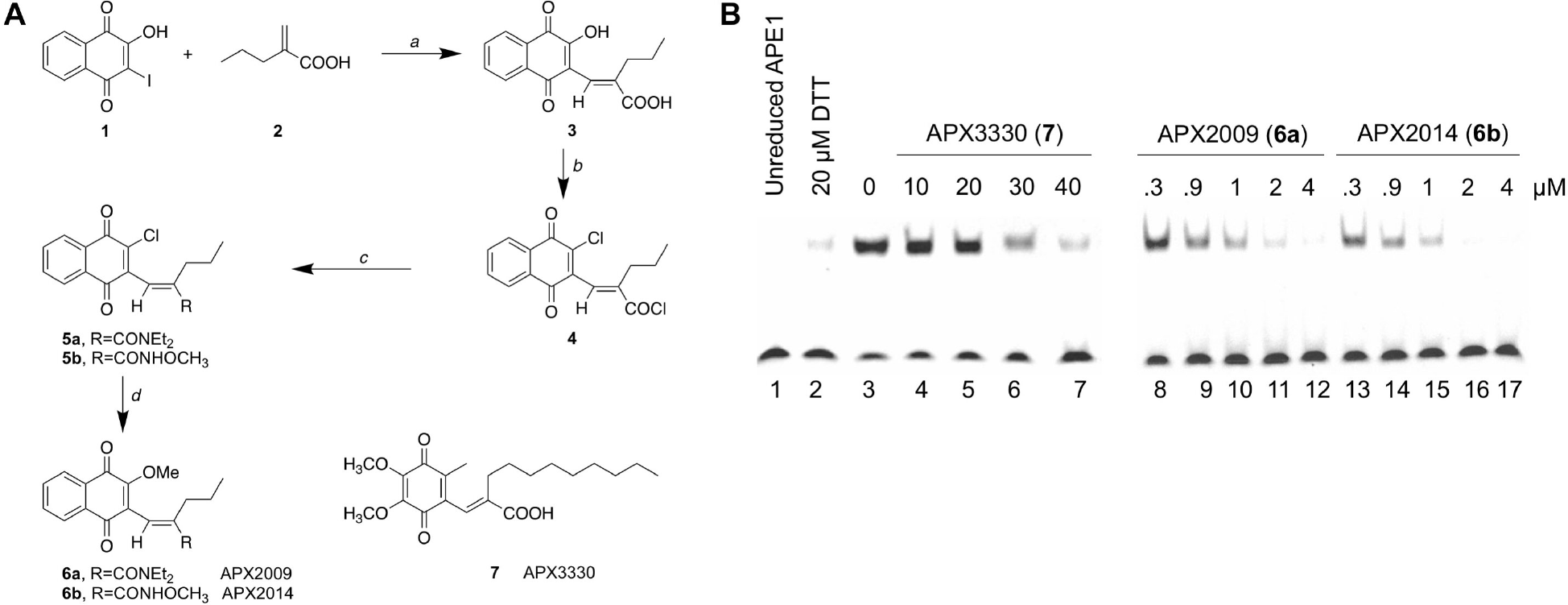
Synthesis and activity of Ref-1 inhibitors. (A) Synthetic scheme for APX2009 (**6a**) and APX2014 (**6b**). Structure of APX3330 (**7**) included for reference. Reagents and conditions: *a*, 2-iodo-3-hydroxy-1,4-naphthoquinone (iodolawsone, **1**), 2-propylacrylic acid (**2**), K_2_CO_3_, Pd(OAc)_2_, argon, 100°C, 1 hour, 74%; *b*, (COCl)_2_, DMF, DCM, RT overnight, 100%; *c*, DEA·HCl (APX2009) or CH_3_ONH_2_·HCl (APX2014), DIPA·HCl, RT, 45 min, 62% and 71% respectively; *d*, NaOCH_3_/CH_3_OH, argon, 30 min, RT, 96% and 86%, respectively. See Supplemental Methods for full synthetic details. (B) APX2009 and APX2014 are more effective inhibitors of Ref-1-induced AP-1 DNA binding than APX3330 in an EMSA. Two separate gels from the same experiment are shown. The IC_50_ for redox EMSA inhibition was 25, 0.45 and 0.2 μM for APX3330, APX2009 and APX2014, respectively. These assays were performed multiple times with similar results.

### Electrophoretic mobility shift assays (EMSA)

These assays were performed as previously described (Luo et al., 2008; Kelley et al., 2011; Su et al., 2011; Luo et al., 2012; Zhang et al., 2013). Briefly, an increasing amount of APX3330, APX2009 or APX2014 was preincubated with purified Ref-1 protein in EMSA reaction buffer for 30 min. The EMSA assay was performed using the AP-1 target DNA sequence and AP-1 protein.

### Cells

Primary human retinal microvascular endothelial cells (HRECs) were obtained from Cell Systems, Inc. (Kirkland, WA), while the Rf/6a macacque choroidal endothelial cell line was obtained from ATCC (Manassas, VA). Cells were maintained as described (Basavarajappa et al., 2017), re-ordered at least annually, and regularly assessed for mycoplasma contamination.

### *In vitro* cell proliferation assay

Endothelial cell proliferation was assessed as described previously (Basavarajappa et al., 2014; Basavarajappa et al., 2017). Briefly, 2.5×10^3^ cells were seeded in 100 μL of growth medium and plated in each well of 96-well clear-bottom black plates and incubated for 24 h. APX2009, APX2014, or DMSO vehicle (DMSO final concentration = 1%) was added, and the plates were incubated for 24-48 h at 37°C and 5% CO2. AlamarBlue reagent (11.1 μL) was added to each well of the plate and 4 h later fluorescence readings were taken at excitation and emission wavelengths of 560 nm and 590 nm, respectively, using a Synergy H1 plate reader (BioTek, Winooski, VT). GI50 was calculated using GraphPad Prism v. 7.0.

### EdU incorporation, Ki-67 Staining, and TUNEL

These assays were carried out as described previously (Basavarajappa et al., 2014; Basavarajappa et al., 2017) but using chamber slides not coverslips. Briefly, cells (30,000 per well) were seeded on 8-well chamber slides coated with attachment factors and allowed to attach overnight. Cells were treated with the indicated compound concentrations for 17 hours (overnight). To assay proliferation, cells were incubated with EdU in complete media for 8 hours at 37°C. Cells were then fixed in 4% paraformaldehyde for 20 min and permeabilized using 0.25% Triton X-100 prepared in PBS. Cells were incubated with a rabbit-specific monoclonal antibody against Ki-67 (D3B5) (#9129; Cell Signaling, Danvers, MA) (1:400) overnight at 4°C. Secondary antibody was Alexa Fluor goat anti-rabbit 488 (A11034; Invitrogen, Carlsbad, CA) with DAPI counter-stain for nuclear staining. Proliferating cells that incorporated EdU were detected using the Click-iT EdU Imaging kit (Invitrogen, Carlsbad, CA). Alternatively, apoptotic cells were visualized using the Click-iT TUNEL assay kit (Invitrogen, Carlsbad, CA) as per the manufacturer’s instructions, with Hoechst 33342 counter-stain for nuclear staining, and a 17-hour treatment with 1 μM staurosporine as positive control. The cells were imaged using a Zeiss AxioImager D2 microscope or an LSM 700 confocal microscope and the percentage of positive cells was counted on three low-power (for TUNEL) or high-power (for Ki-67 and EdU) fields per well using ImageJ software.

### Cell cycle analysis

HRECs (2×10^6^) were grown in EGM-2 medium. Cells were serum starved in EBM-2 medium overnight, then treated with the indicated concentrations of APX2009 or APX2014 along with DMSO control for 24 hours in complete medium. Cells were washed twice in ice-cold PBS followed by fixation in 66% ethanol solution overnight at 4°C. Fixed cells were again washed twice in ice-cold PBS and the pellets were resuspended in propidium iodide staining solution for 30 mins at 37°C (20 μg/mL propidium iodide prepared in 1× PBS containing 0.1% Triton X-100 and 100 μg/mL RNase A). After incubation cells were analyzed using flow cytometry (FACSCalibur, BD Biosciences, San Jose, CA). Pulse shape analysis was used to exclude doublets and debris. The single cell population was then assessed by the FL2 area histogram plot using ModFit software (v. 5.0) and cell cycle profiles were generated.

### *In vitro* cell migration assay

Endothelial cell migration was monitored as described before (Basavarajappa et al., 2014; Basavarajappa et al., 2017). Briefly, HRECs and Rf/6a were grown until confluency in 12-well plates. Using a sterile 10-μL micropipette tip, a scratch wound was made across the center of each well and fresh complete media containing DMSO or different concentrations of APX2009 or APX2014 compounds were added to the wells (DMSO final concentration = 1%). Wells were imaged via digital brightfield microscopy at different time points, and the number of migrated cells into the scratched area was manually counted.

### *In vitro* Matrigel tube formation assay

The ability of HRECs and Rf/6a cells to form tubes *in vitro* was monitored as described before (Basavarajappa et al., 2014; Basavarajappa et al., 2017). Briefly, 1.5×10^4^ cells in 100 μL of growth medium containing DMSO or APX compounds were added to each well of a 96-well plate that was pre-coated with 50 μL per well of Matrigel basement membrane (DMSO final concentration = 1%). Brightfield digital micrographs of each well at different time points were taken to measure the *in vitro* tube formation using the Angiogenesis Analyzer plugin in ImageJ software (v.1.48; http://image.bio.methods.free.fr/ImageJ/?Angiogenesis-Analyzer-for-ImageJ.html).

### NF-κB p65 nuclear translocation assay

The NF-κB nuclear translocation assay was performed by seeding 30,000 HRECs/well on an 8-well chamber slide coated with attachment factors. The cells were grown in EGM-2 medium overnight before treating with indicated concentrations of APX2009 and APX2014, or 10 μM BAY 11-7082 (Sigma, St. Louis, MO) as a positive control NF-κB inhibitor. After 17 h incubation, the media was replaced with EBM-2 (minimal medium) with indicated concentrations of compound or DMSO for 1 hour. The cells were then stimulated with 10 ng/ml TNF-α in EBM-2 for 20 min at 37°C to activate NF-κB. Cells were then fixed in 4% paraformaldehyde and permeabilized using 0.5% Triton X-100 solution prepared in PBS. The cells were incubated with a monoclonal antibody against NF-κB p65 (sc-8008; Santa Cruz, Santa Cruz, CA) (1:50) overnight at 4°C, followed by Alexafluor 555 goat anti-mouse secondary antibody (1:2000) for one hour. The cells were counter-stained with Hoechst 33342 for nuclear staining and then mounted using Everbrite hardset mounting medium. The cells were imaged using a Zeiss AxioImager D2 microscope.

### qRT-PCR

The assay was performed as described previously (Basavarajappa et al., 2014; Basavarajappa et al., 2017). RNA was extracted from cells treated as indicated using Trizol (Invitrogen). cDNA was synthesized from 1 μg RNA using random primers and iScript reverse transcriptase (Bio-Rad, Hercules, CA). qPCR was performed in 10 μL volumes in a 384-well plate, with Fast Advanced Master Mix and TaqMan probes on a ViiA7 thermal cycler (Applied Biosystems, Foster City, CA).

Primer/probesets used were as follows: *VEGFA* (Hs00900055_m1), *VCAM1* (Hs01003372_m1), and *CCL20* (Hs01011368_m1), and housekeeping controls *HPRT* (Hs02800695_m1) and *TBP* (Hs00427620_m1). The data were analyzed using the *ΔΔC*_*t*_ method. The expression levels of genes were normalized to the two housekeeping genes and calibrated to the DMSO treated sample.

### Animals

All animal experiments were approved by the Indiana University School of Medicine Institutional Animal Care and Use Committee and followed the guidelines of the Association for Research in Vision and Ophthalmology Statement for the Use of Animals in Ophthalmic and Visual Research. Wild-type female C57BL/6 mice, 6-8 weeks of age, were purchased from Envigo (Indianapolis, IN; for choroidal sprouting experiments) or Jackson Laboratory (Bar Harbor, ME; for L-CNV) and housed under standard conditions (Wenzel et al., 2015). Mice were anesthetized for all procedures by intraperitoneal injections of 90 mg/kg ketamine hydrochloride and 5 mg/kg xylazine, with intraperitoneal atipamezole reversal (1 mg/kg). Treatments were randomly assigned by cage.

### Choroidal sprouting assay

*Ex vivo* Choroidal sprouting was assessed as described previsouly (Sulaiman et al., 2016; Basavarajappa et al., 2017). Briefly, choroid-sclera was dissected from 7 to 8 week old mouse eyes and pieces were embedded in Matrigel (growth factor reduced) and grown in EGM-2 medium containing antibiotics for 72 h to allow sprouting to initiate. The indicated concentrations of APX2009 and APX2014 compounds (in DMSO, final DMSO concentration 0.5 and 0.2%, respectively) were added and growth allowed to proceed for 48 h. Images were taken and growth was quantified by measuring the distance from the edge of the choroidal piece to the growth front in four directions per sample using ImageJ software.

### Laser-induced choroidal neovascularization

L-CNV was induced as described previously (Sulaiman et al., 2015; Sulaiman et al., 2016; Basavarajappa et al., 2017). Studies were powered to have an 80% chance of detecting effect size differences of 50%, assuming 30% variability, α = 0.05. Briefly, pupils of anesthetized mice were dilated with 1% tropicamide (Alcon Laboratories Inc., Forth Worth, TX) and lubricated with hypromellose ophthalmic demulcent solution (Gonak) (Akorn, Lake Forest, IL). A coverslip was used to allow viewing of the posterior pole of the eye. Three burns of a 532 nm ophthalmic argon green laser coupled with a slit lamp (50 μm spot size, 50 ms duration, and 250 mW pulses) were delivered to each 3, 9, and 12 o’clock position, two-disc diameters from optic disc. The bubbling or pop sensed after laser photocoagulation was considered as the successful rupture of Bruch’s membrane. Lesions in which bubbles were not observed were excluded from the study. To assess the antiangiogenic activity of APX3330, the mice were i.p. injected with compound (50 mg/kg body weight), twice daily, five days on/two days off, as used previously *in vivo* (Fishel et al., 2011; Lou et al., 2014; Biswas et al., 2015). Vehicle was 4% Cremophor:ethanol (1:1) in PBS. For APX2009, doses were 12.5 mg/kg or 25 mg/kg body weight, twice daily until 14 days of laser treatment unless otherwise indicated. Vehicle was propylene glycol, Kolliphor HS15, Tween 80 (PKT) (McIlwain et al., 2018). Mice were weighed daily.

### *In vivo* imaging

Optical coherence tomography (OCT) was performed in L-CNV mice as described previously (Sulaiman et al., 2016), at the indicated times using the Micron III intraocular imaging system (Phoenix Research Labs, Pleasanton, CA). Briefly, before the procedure, eyes of anesthetized mice were dilated with 1% tropicamide solution (Alcon, Fort Worth, TX) and lubricated with hypromellose ophthalmic demulcent solution (Gonak) (Akorn, Lake Forest, IL, USA). Mice were then placed on a custom heated stage that moves freely to position the mouse eye for imaging. Several horizontal and vertical OCT images were taken per lesion. Fluorescein angiography was performed 14 days post laser by intraperitoneal injection of 50 μL of 25% fluorescein sodium (Fisher Scientific, Pittsburgh, PA). Fundus images were taken using the Micron III system and Streampix software.

### Choroidal flatmount immunofluorescence

Mouse eyes were harvested 14 days after L-CNV induction. The eyes were enucleated and fixed in 4% paraformaldehyde/PBS overnight. The anterior segment, lens, and retina were removed, and the posterior eye cups were prepared for choroidal flat mounts. The posterior eye cups were washed with PBS and permeabilized in blocking buffer containing 0.3% Triton X-100, 5% bovine serum albumin (BSA) in PBS for two hours at 4°C. The eye cups were then double stained for vasculature with rhodamine-labeled *Ricinus communis* agglutinin I (Vector Labs, Burlingame, CA) and Alexa Fluor^TM^ 488 conjugated-Isolectin B4 from *Griffonia simplicifolia* (GS-IB4) (Molecular Probes, Thermo Fisher Scientific) at 1:250 dilution in buffer containing 0.3% Triton X-100, 0.5% BSA in PBS for 16-20 hours at 4°C. After antibody incubation, whole mounts were washed three times with PBS for 15 mins each step at 4°C with 0.1% Triton X-100. After washing, choroidal flatmounts were mounted in aqueous mounting medium (VectaShield; Vector Laboratories, Inc.) and coverslipped for observation by confocal *Z*-stack imaging (LSM 700, Zeiss, Thornwood, NY) to estimate lesion volume. The sum of the stained area in each optical section, multiplied by the distance between sections (3 μm), gave the CNV lesion volume and lesion volume was quantified using ImageJ software. Lesions were only included for analysis if they met quality control standards as published (Poor et al., 2014). All lesions in an eye were averaged to represent a single *n*.

### Statistical analyses

Statistical analyses were performed with GraphPad Prism 7 software. One-way ANOVA was used with Dunnett’s post hoc test for EdU, Ki-67, TUNEL, cell cycle, migration, tube formation, qPCR, and choroidal sprouting experiments. Unpaired Student’s *t*-test was used for the APX3330 *in vivo* experiment. One-way ANOVA was used with Tukey’s post hoc test for L-CNV analysis in APX2009 *in vivo* experiments, while repeated measures two-way ANOVA was used to compare body weights between treatments and over time. Two-sided *p* values < 0.05 were considered statistically significant.

## Results

### Novel Ref-1 inhibitors are more potent than APX3330 in blocking binding of AP-1 DNA binding

We synthesized APX2009 (**6a**) and APX2014 (**6b**) (Fig. 1A) and demonstrated that both compounds had enhanced inhibition of Ref-1-induced transcription factor binding to DNA compared to APX3330 (**7**) (Fig. 1B), while having substantially different physiochemical properties. The new compounds have lower molecular weights, and lack the carboxylate group and long alkyl chain of APX3330. The new compounds also have significantly reduced lipophilicity as determined by computer based calculation of their clogP values, APX3330 = 4.5, APX2009 = 2.7, and APX2014 = 1.9.

### APX2009 and APX2014 block endothelial cell proliferation

Endothelial cell proliferation with increased survival supports the cells that make up new blood vessels, leading to angiogenesis. As an initial test of the antiangiogenic potential of our novel Ref-1 inhibitors, we assessed their ability to inhibit the proliferation of HRECs and Rf/6a choroidal endothelial cells (Fig. 2). Both compounds dose-dependently blocked proliferation of both cell types in an alamarBlue assay, with APX2014 5− to 10-fold more potent than APX2009. Primary HRECs were more sensitive to both compounds than the Rf/6a choroidal cell line, as seen with other antiangiogenic compounds (Basavarajappa et al., 2017).

**Fig. 2.**
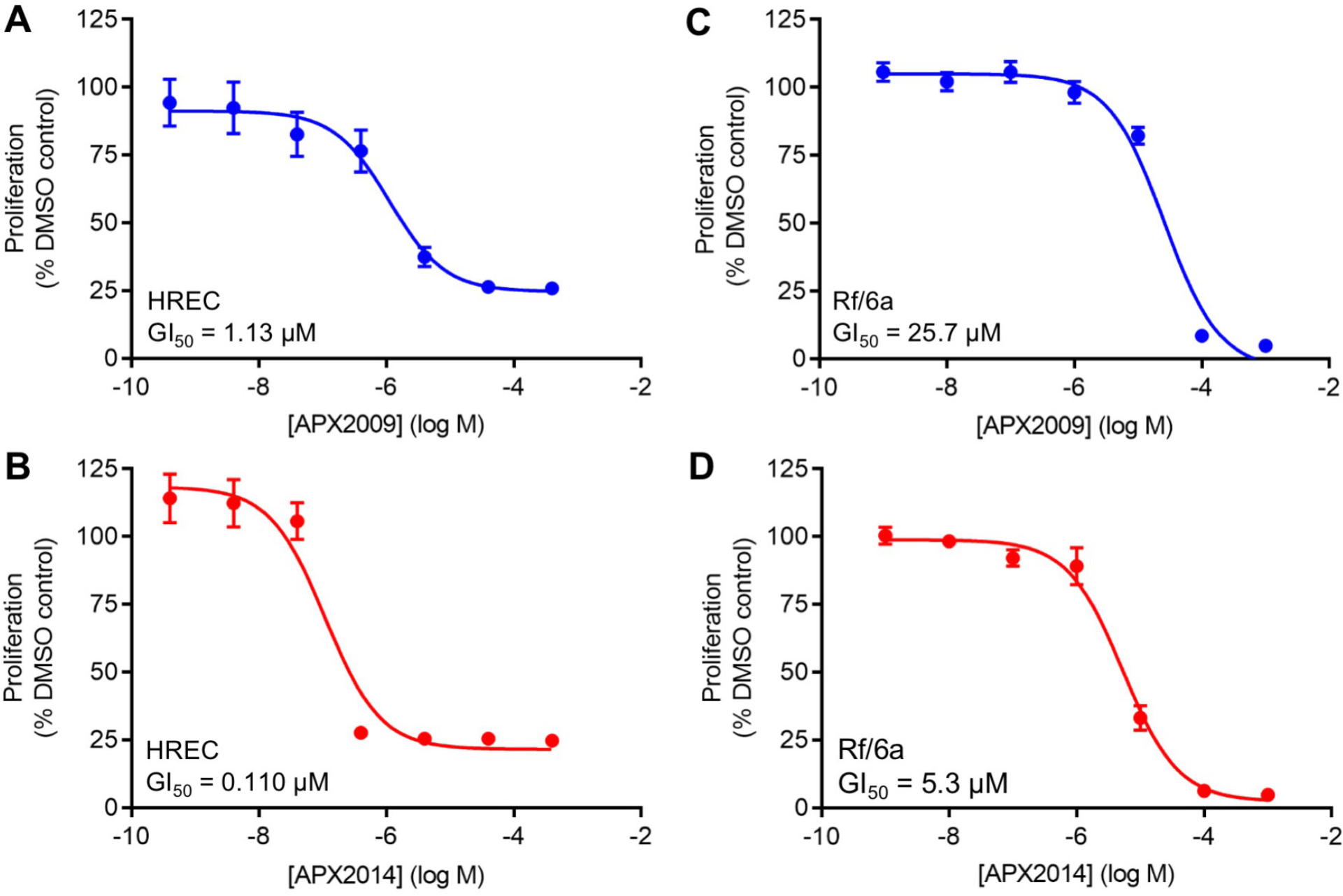
Compounds APX2009 and APX2014 inhibit endothelial cell proliferation in HRECs and Rf/6a cells *in vitro*. Dose dependent effects of APX2009 (A) and APX2014 (B) in human retinal endothelial cells (HRECs), and dose dependent effect of APX2009 (C) and APX2014 (D) in Rf/6a choroidal endothelial cells. *In vitro* proliferation was measured using an alamarBlue assay. Median growth inhibition (GI50) values are indicated. Mean ± S.E.M., *n* = 3 per dose.

### APX2009 and APX2014 block S phase without inducing apoptosis

We assessed the activity of our novel compounds in more detail in HRECs. Both compounds reduced the number of cells going through S phase as evidenced by reduced Ki-67 staining and reduced EdU incorporation (Fig. 3; Supplemental Figs. 1A & 2). This was also evident as a modest increase in cells in G0/G1 phase at high doses of compound, with a concomitant decrease in G2/M phase cells (Supplemental Fig. 1B, C). However, neither compound induced apoptosis at anti-proliferative doses as assessed by TUNEL (Supplemental Fig. 3).

**Fig. 3.**
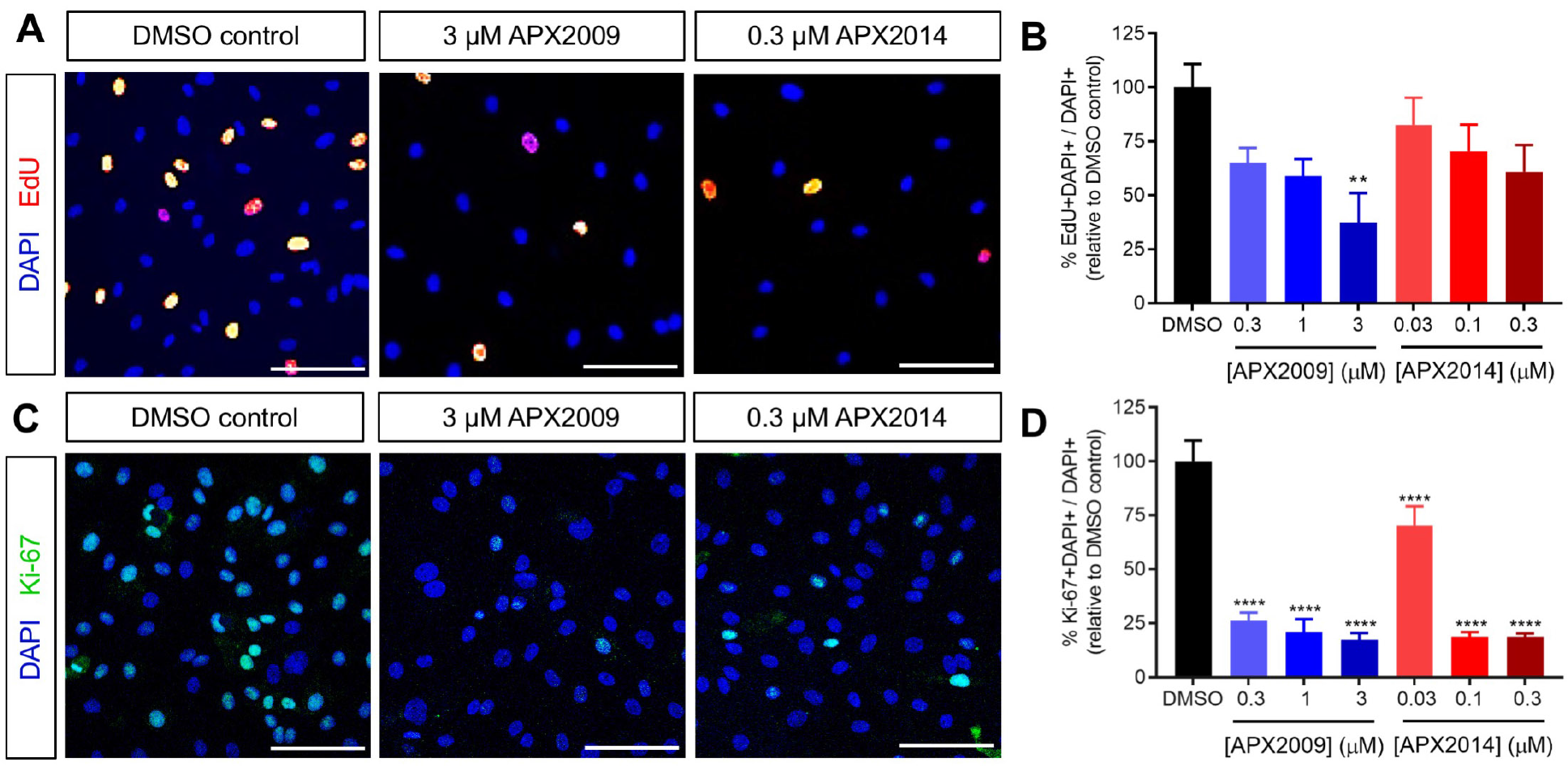
Compounds APX2009 and APX2014 inhibit S phase in HRECs. After treating HRECs with the indicated concentrations of APX2009 and APX2014, (A) EdU (red) and (C) Ki-67 (green) were detected and nuclei (blue) stained with DAPI; Scale bar = 100 μm. (B) Quantification of EdU and (D) quantification of Ki-67 in HRECs. Mean ± S.E.M., *n* = 3 fields per dose. **, *p* < 0.01; ****, *p* < 0.0001 compared to DMSO control (one-way ANOVA with Dunnett’s post hoc test). Representative data from three independent experiments. See Supplemental Figs. 1A & 2.

### APX2009 and APX2014 block endothelial cell migration

Neovascularization involves an array of coordinated events, including extracellular matrix degradation, cell migration, cell proliferation, and morphogenesis of endothelial cells. To know the effect of APX2009 and APX2014 compounds on endothelial cell migration, a scratch-wound assay was performed. (Fig. 4; Supplemental Figs. 4 & 5). Both compounds again were dose-dependently and significantly effective here, without causing obvious cytotoxicity over the short time course of these assays.

**Fig. 4.**
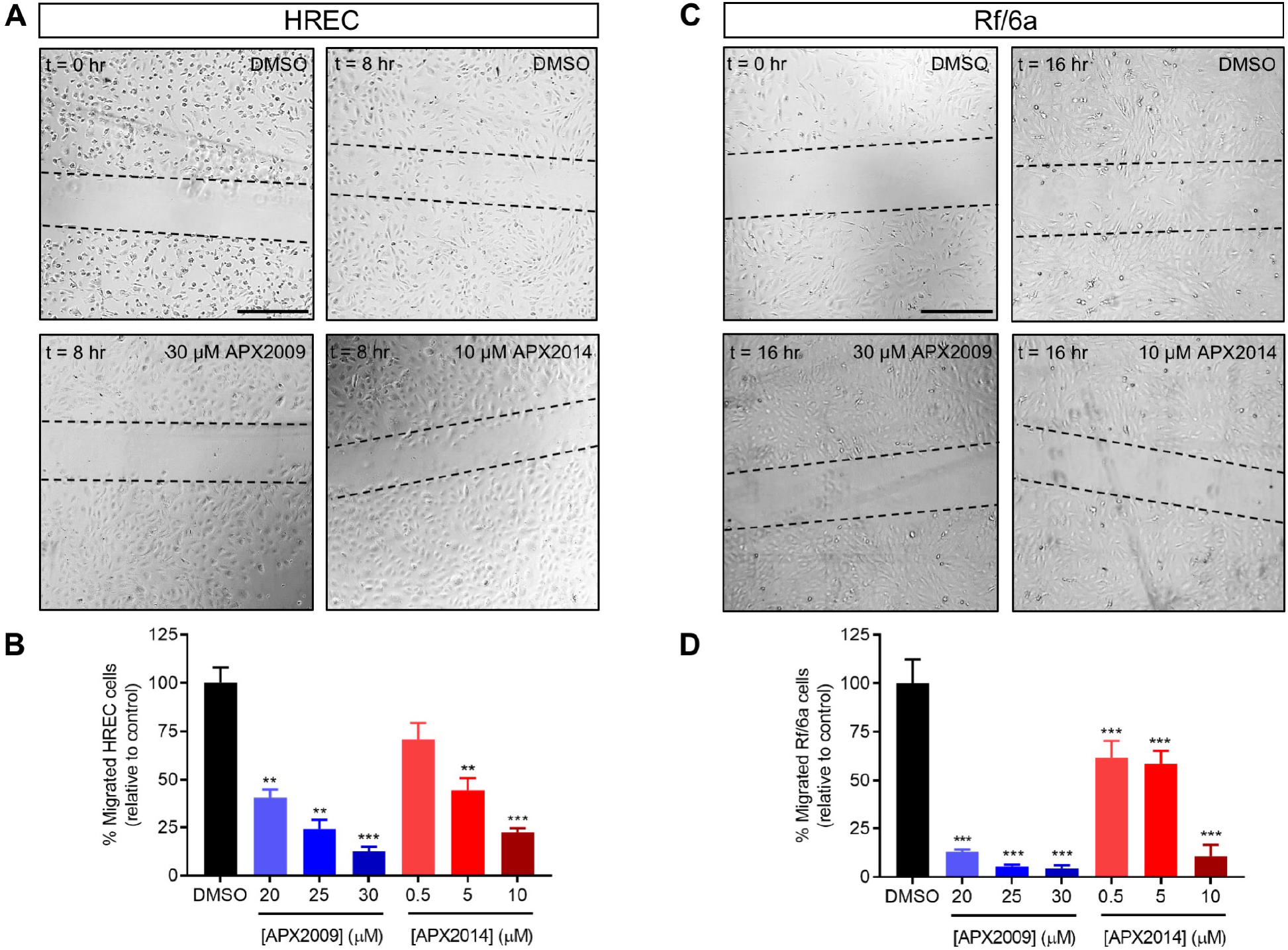
Compounds APX2009 and APX2014 inhibit endothelial cell migration in HRECs and Rf/6a cells *in vitro*. (A) Effect of APX2009 and APX2014 on cell migration in HRECs. A confluent monolayer of HRECs with various treatments (highest doses shown) was wounded and wound closure was monitored for 8 hours. (B) Quantitative analysis of cell migration shows that APX compounds significantly block the migration of HRECs. (C) Effect of APX2009 and APX2014 on cell migration in Rf/6a cells. A confluent monolayer of Rf/6a with various treatments (highest doses shown) was wounded and wound closure was monitored for 16 hours. (D) Quantitative analysis of cell migration shows that APX compounds significantly block the migration of Rf/6a cells. Mean ± S.E.M., *n* = 3 per dose. **, *p* < 0.01; ***, *p* < 0.001 compared to DMSO control (one-way ANOVA with Dunnett’s post hoc test). Representative data from three independent experiments. Scale bar = 500 μm. See Supplemental Figs. 4 & 5.

### APX2009 and APX2014 block endothelial cell tube formation

Endothelial cells organize and form capillary-like structures upon plating on an extracellular matrix such as Matrigel. The organization of endothelial cells into a three-dimensional network of tubes is essential for angiogenesis. As such, the Matrigel tube formation assay is a good *in vitro* predictor of angiogenic potential *in vivo*. In this assay, both APX2009 and APX2014 inhibited tubule formation markedly, at concentrations lower than those required for inhibiting migration alone, strongly indicative of antiangiogenic activity (Fig. 5; Supplemental Figs. 6 & 7).

**Fig. 5.**
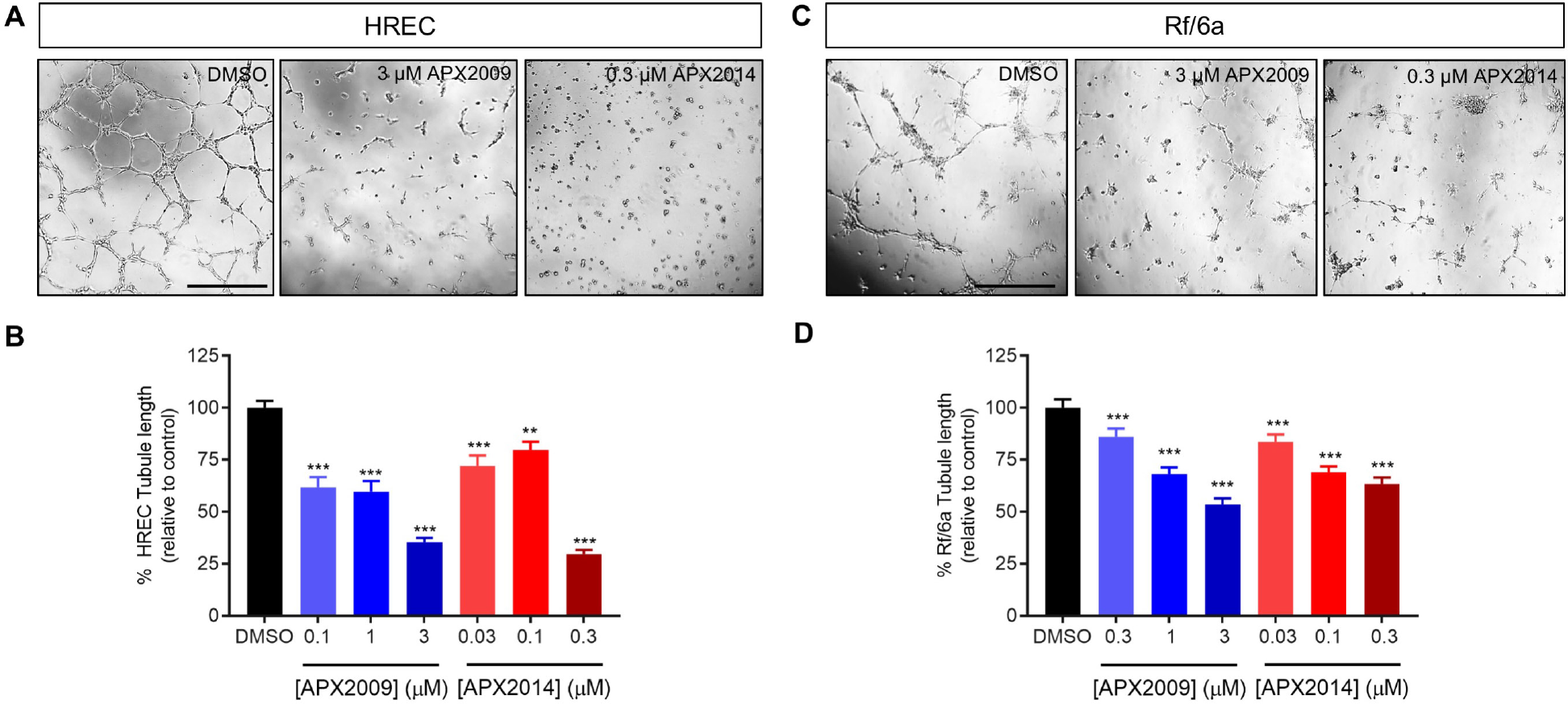
Compounds APX2009 and APX2014 inhibit endothelial tube formation in HRECs and Rf/6a cells *in vitro*. Representative images of tube formation on Matrigel by HRECs (A) and Rf/6a cells (C) in the presence of the indicated concentrations of APX compounds; Quantitative analysis of tube formation in HRECs (B) and Rf/6a cells (D) following APX2009 and APX2014 treatment. Tubular length was measured and represented as relative to DMSO control. Mean ± S.E.M., *n* = 3 wells. **, *p* < 0.01; ***, *p* < 0.001 compared to DMSO control (one-way ANOVA with Dunnett’s post hoc test). Representative data from three independent experiments. Scale bar = 500 μm. See Supplemental Figs. 6 & 7.

### APX2009 and APX2014 inhibit NF-κB activity

Since Ref-1 inhibition has previously been associated with reduction in NF-κB activity (Shah et al., 2017), we assessed activity of this pathway in response to our novel compounds in HRECs, to determine if APX2009 and APX2014 were acting through the expected mechanisms.

First, we assayed translocation of the p65 subunit of NF-κB into the nucleus in response to TNF-α, a key indication of pathway activity. Translocation of p65 was dose-dependently attenuated in APX2009 and APX2014-treated HRECs (Fig. 6A). Moreover, production of the mRNA of *VEGFA*, *VCAM1*, and *CCL20*, all of which are downstream of NF-κB, was decreased 3-to 10-fold by these compounds (Fig. 6B, C, D).

**Fig. 6.**
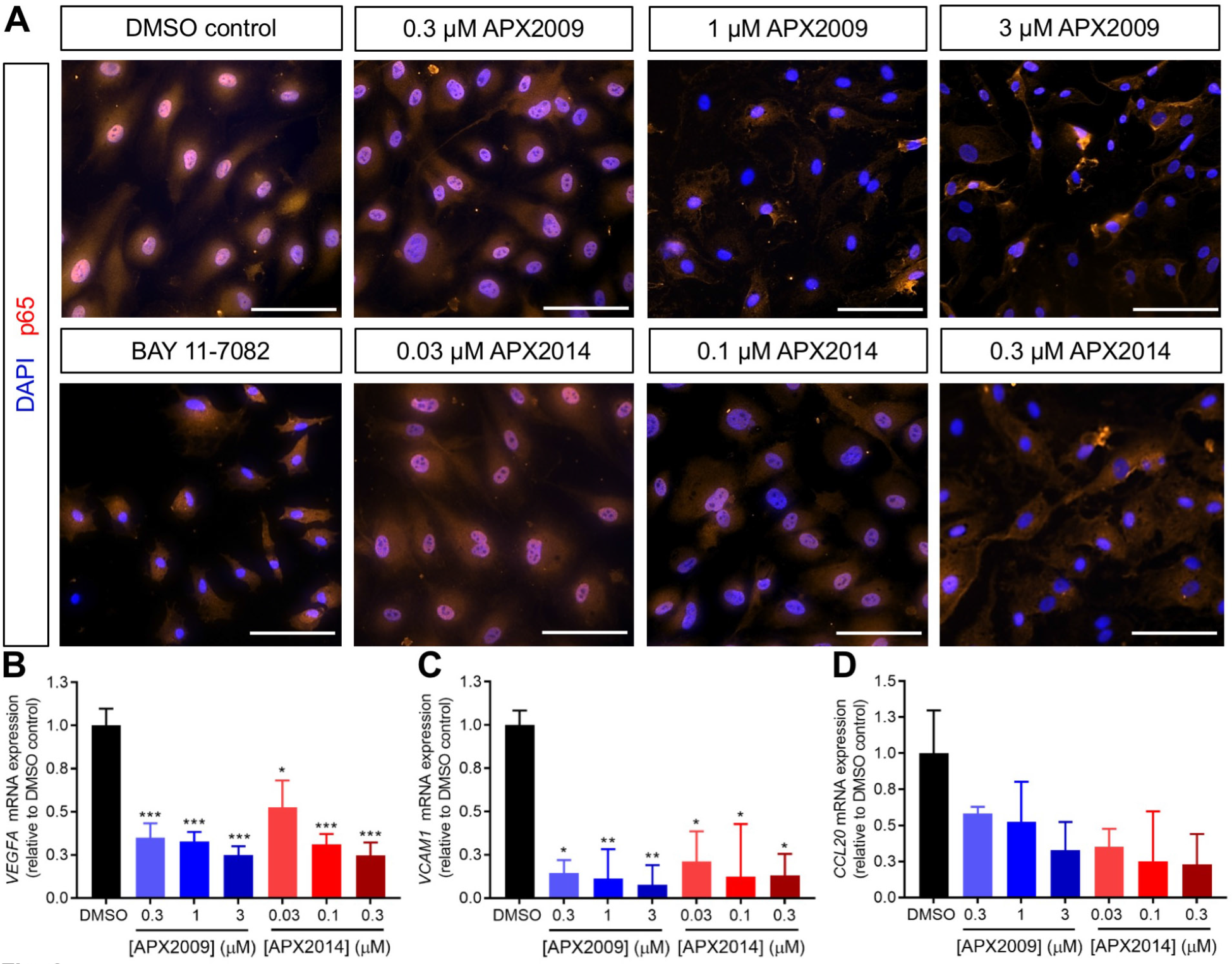
Compounds APX2009 and APX2014 inhibit TNF-α mediated NF-κB signaling and proangiogenic target gene mRNA expression. After treating HRECs with the indicated concentrations of APX2009 and APX2014, p65 (red) was detected by immunofluorescence and nuclei (blue) stained with DAPI; compounds dose-dependently reduced p65 nuclear translocation as evidenced by decreased overlap between red and blue signals. BAY 11-7082 is a positive control NF-κB inhibitor. Scale bar = 100 μm. (B) *VEGFA*, (C) *VCAM1*, and (D) *CCL20* mRNA expression levels in HRECs. APX2009 and APX2014 dose dependently inhibited levels of each transcript. Mean ± S.E.M., *n* = 3 technical replicates. *, *p* < 0.05; **, *p* < 0.01; ***, *p* < 0.001 compared to DMSO control (one-way ANOVA with Dunnett’s post hoc test). Representative data from three independent experiments.

### APX2009 and APX2014 block angiogenesis *ex vivo*

As a further test of antiangiogenic activity, we employed a choroidal sprouting assay using murine choroidal explants to test the effect of the APX compounds in a complex microvascular bed in tissues (Fig. 7). In this assay, choroidal cells grow out of the choroidal tissue piece into a surrounding Matrigel matrix. Both compounds significantly reduced sprouting, with APX2014 remaining more potent. At 10 μM, APX2009 reduced sprouting by ~70% compared to control (Fig. 7A, B), while at 1 μM (the highest concentration tested), APX2014 reduced sprouting by ~60% compared to control (Fig. 7C, D).

**Fig. 7.**
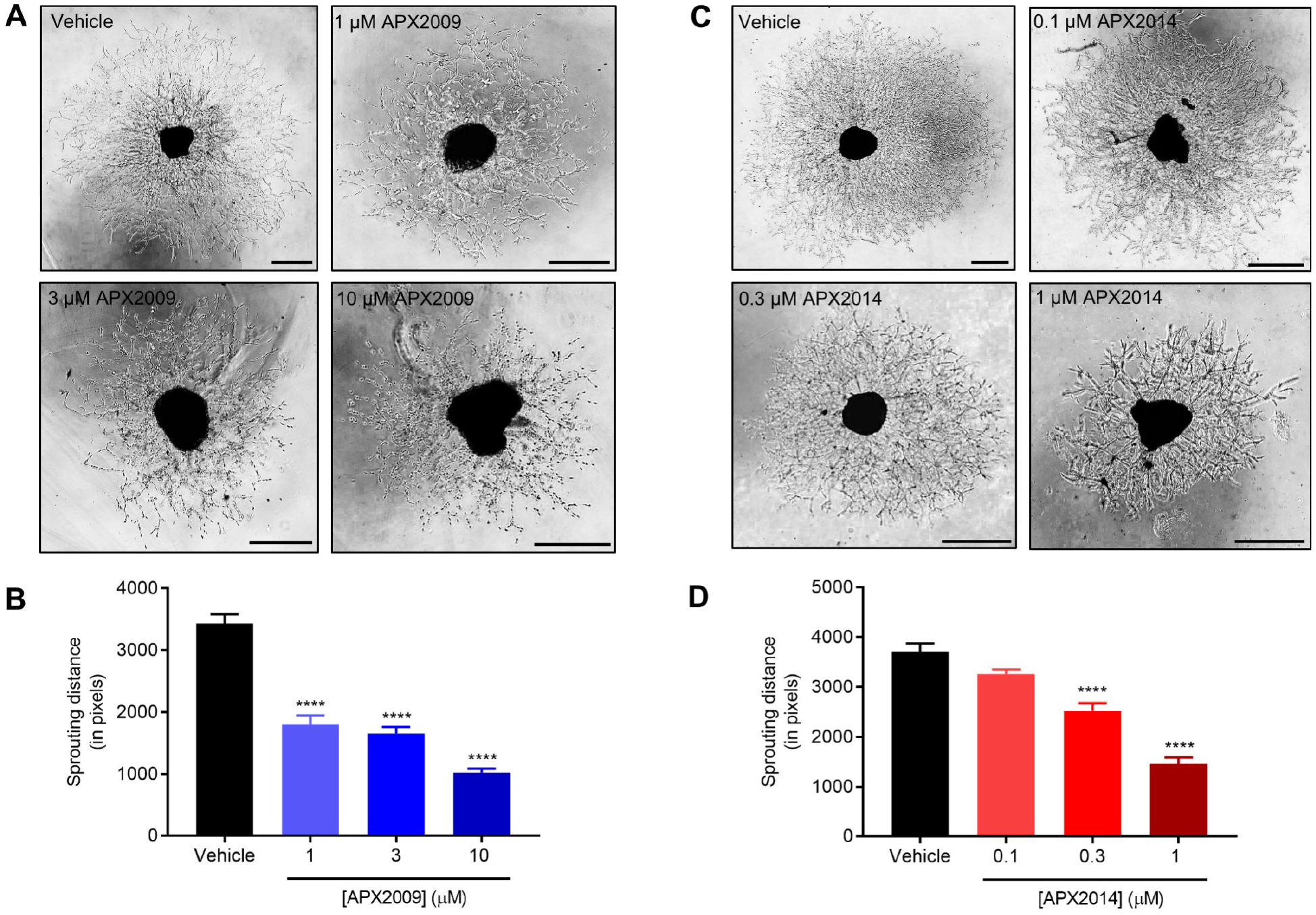
Compounds APX2009 and APX2014 inhibit choroidal sprouting in a concentration-dependent manner. Representative phase contrast images of choroidal sprouts formed 48 hours after treatment with indicated concentrations of APX2009 (A) or APX2014 (C) or vehicle control (0.5% or 0.2% DMSO, respectively). Quantification of sprouting distance from the edge of the APX2009-treated (B) or APX2014-treated (D) choroidal tissue piece to the end of the sprouts averaged from four perpendicular directions using ImageJ software. Mean ± S.E.M., *n* = 4-6 choroids/per treatment; *N* = 3-4 eyes.***, *p* < 0.001; ****, *p* < 0.0001 (ANOVA with Dunnett’s post hoc test). Scale bars = 500 μm.

### Systemic Ref-1 inhibition with parent compound APX3330 can prevent L-CNV

Previous efforts to attenuate ocular neovascularization by Ref-1 inhibition using APX3330 relied on intravitreal delivery of compound (Jiang et al., 2011; Li et al., 2014a; Li et al., 2014b). Although this is the delivery route of the standard-of-care anti-VEGF biologics and ensures that the drug gets to the right place, in humans it is labor-intensive, causes patient discomfort, and incurs a risk of potentially vision-threatening endophthalmitis (Day et al., 2011). Thus, we explored if systemic (intraperitoneal) administration of Ref-1 inhibitors could offer an alternative route of therapy for L-CNV. As a proof-of-concept, we tested i.p. injections of the first-generation Ref-1 inhibitor APX3330 (**7**) delivered 50 mg/kg twice daily, 5 days on/two days off, for two weeks. This dosing regimen was chosen as it was previously successful and non-toxic for preclinical tumor studies (Fishel et al., 2011; Lou et al., 2014; Biswas et al., 2015). Animals treated with APX3330 displayed significantly reduced L-CNV volume (Fig. 8).

**Fig. 8.**
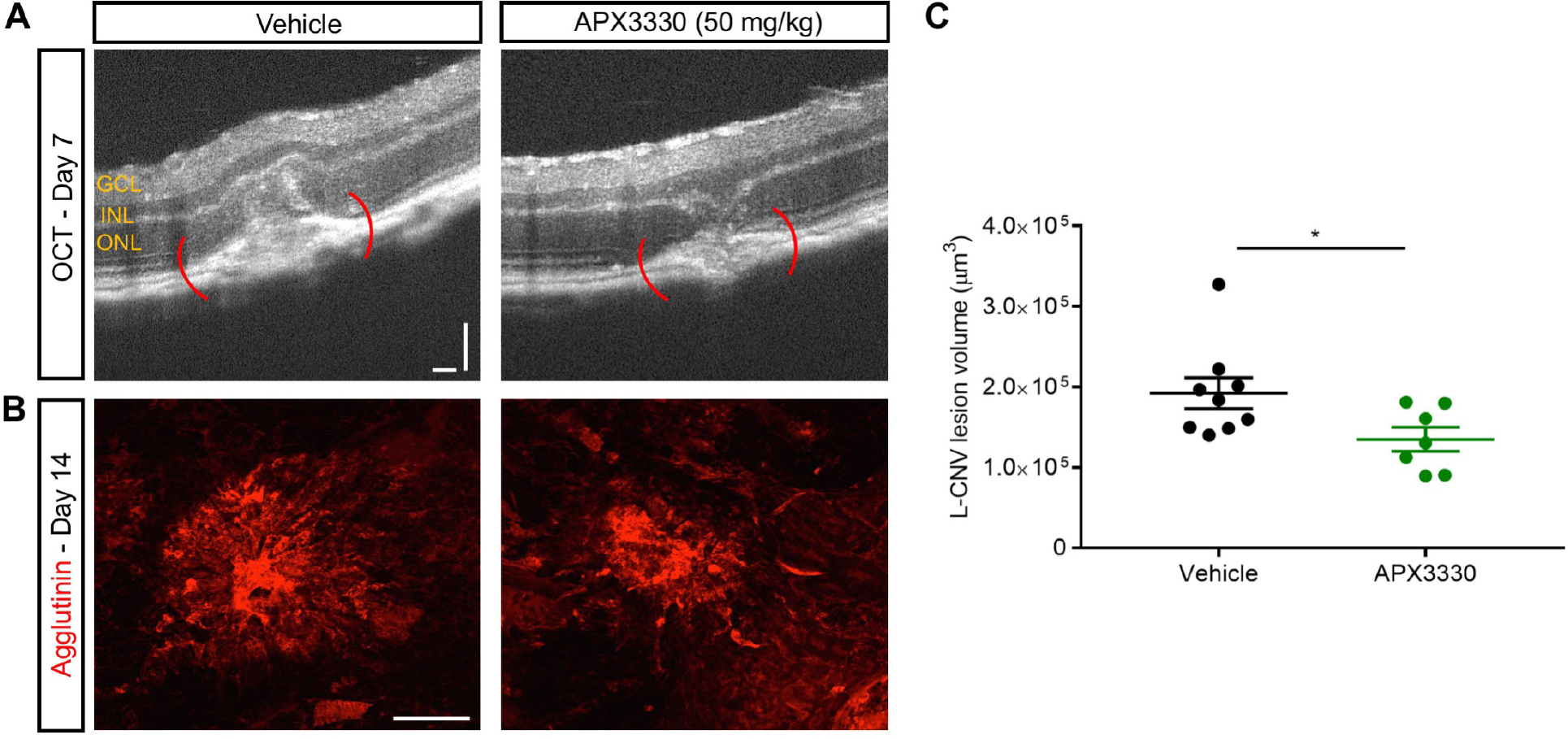
Systemic Ref-1 inhibition with APX3330 blocks neovascularization in the laser-induced choroidal neovascularization (L-CNV) mouse model. (A) Representative optical coherence tomography (OCT) images obtained 7 days post-laser, showing CNV lesions in eyes of vehicle (left) and 50 mg/kg i.p. APX3330 treated animals (right). (B) Representative images from confocal microscopy for agglutinin-stained CNV lesions 14 days post-laser treatment. (C) Quantification of CNV lesion vascular volumes from *Z*-stack images at day 14 using ImageJ software. Mean ± S.E.M., *n* = 7–9 eyes/treatment. * *p* < 0.05 (unpaired Student’s *t*-test). Scale bars = 100 μm. GCL, ganglion cell layer; INL, inner nuclear layer; ONL, outer nuclear layer.

### Systemic administration of more potent derivative APX2009 reduces L-CNV significantly

Given that APX3330 reduced L-CNV with systemic adminstration, we explored if similar effects could be observed with our second-generation Ref-1 inhibitors. We chose APX2009 for this experiment as it had previously been safely dosed in animals (Kelley et al., 2016). We used two dosage regimens previously employed, 12.5 or 25 mg/kg twice daily for two weeks. The lower dose did not reduce L-CNV, but the 25 mg/kg dose had a marked effect (Fig. 9). This was qualitatively evident by OCT imaging on Day 7, and even more substantial on Day 14 (Fig. 9A). In addition, reduced fluorescein leakage was seen in lesions by fluorescein angiography at Day 14 (Fig. 9B). Finally, L-CNV lesion volume assessed by *ex vivo* staining with agglutinin (Fig. 9C) and isolectin B4 (Supplemental Fig. 8), was reduced by 25 mg/kg APX2009 approximately four-fold compared to vehicle (Fig. 9D).

**Fig. 9.**
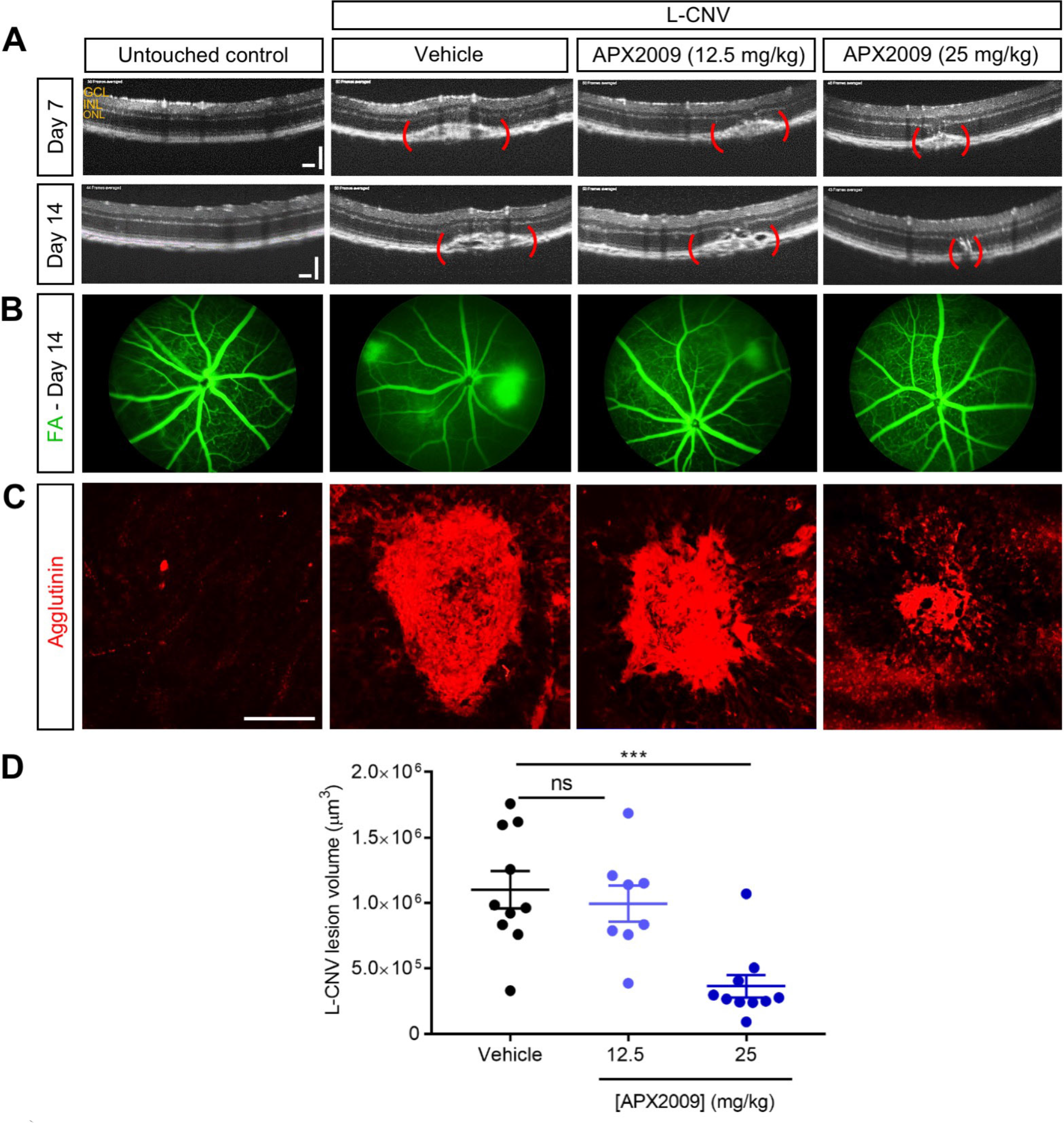
Intraperitoneal APX2009 inhibits choroidal neovascularization in the L-CNV mouse model. (A) Representative OCT images obtained 7 and 14 days post-laser, showing CNV lesions of untouched control, vehicle, 12.5 mg/kg and 25 mg/kg APX2009 compound i.p. injected twice daily until 14 days post-laser treatment. (B) Fluorescein angiography (FA) of CNV showing the vascular leakage suppression by APX2009. (C) Representative images from confocal microscopy for agglutinin-stained CNV lesions 14 days post-laser treatment. (D) Quantification of CNV lesion vascular volumes from *Z*-stack images at day 14 using ImageJ software. Mean ± S.E.M., *n* = 8- 10 eyes/treatment. ns, non-significant; ***, *p* < 0.001 compared to DMSO control (one-way ANOVA with Tukey’s post hoc test). Scale bars = 100 μm. GCL, ganglion cell layer; INL, inner nuclear layer; ONL, outer nuclear layer. See also Supplemental Fig. 5.

## Discussion

We have identified two new Ref-1 inhibitors, APX2009 and APX2014, and demonstrated antiangiogenic activity of these compounds. Moreover, we showed for the first time that *systemic administration* of Ref-1 inhibitors can attenuate L-CNV. As L-CNV is a widely-used model that recapitulates the choroidal neovascularization underlying wet AMD, Ref-1 inhibition could find therapeutic utility for this indication. Our *in vitro* data, and previous work (Jiang et al., 2011), suggest that Ref-1 inhibition also effectively blocks angiogenesis involving retinal endothelial cells. Thus, these inhibitors may also be useful for retinal neovascular diseases like ROP and PDR.

The observed effects are likely attributable to inhibition of Ref-1 redox signaling, rather than DNA repair inhibition, as the compounds are specific for redox signaling inhibition. The molecularly distinct functional portions of Ref-1, redox and DNA repair, are completely independent. For example, mutations of the cysteine at position 65 (C65A) of Ref-1/APE1 abrogate the redox function, but do not affect DNA repair function, and vice versa (McNeill and Wilson, 2007; Vasko et al., 2011). Moreover, Ref-1 inhibitors such as APX3330 do not inhibit APE1 endonuclease activity. In fact, APX3330 and APX2009 can *enhance* APE1 repair activity in neurons (Kelley et al., 2016), potentially contributing to a neuroprotective effect of these agents, which could offer an added benefit in the context of photoreceptor cell death in neovascular eye diseases.

Given their anti-Ref-1 redox signaling activity, APX2009 and APX2014 likely exert their antiangiogenic effects by blocking the activation of transcription factors induced by Ref-1. This includes NF-κB as shown, and also possibly HIF-1α; both of these can regulate VEGF (Forsythe et al., 1996). In retinal pigment epithelial cells, APX3330 reduced both NF-κB and HIF-1α activity, with a concomitant reduction in VEGF expression (Li et al., 2014a; Li et al., 2014b), as we have shown here for APX2009 and APX2014. Additionally, APX3330 treatment of stroke in type one diabetes mellitus rats significantly decreased total vessel density and VEGF expression (Yan et al., 2018). However, the full spectrum of transcription factors modulated by Ref-1 inhibition in the context of ocular neovascularization remain to be determined.

We did not observe obvious intraocular or systemic toxicity of the two compounds tested *in vivo* (APX3330 and APX2009), nor did we see apoptosis or substantial cell death in migration, tube formation, and choroidal sprouting assays. These findings are consistent with the excellent safety profile for APX3330 in humans (Shah et al., 2017). Nonetheless, ocular toxicity of the new compounds and intraocular pharmacokinetics remain to be thoroughly examined.

A well-tolerated, systemic drug therapy has significant potential for treatment of neovascular eye diseases. The existing approved drugs are all biologics requiring intravitreal injection in the context of an ophthalmologist’s clinic. An orally bioavailable drug (as with APX3330) could be administered at home, potentially as a once daily pill. The tradeoff for such a therapy would be much more frequent dosing than that required for intravitreal injections (monthly or less), and more substantial systemic exposure than that seen with intravitreal therapies. But given the strong safety profile of Ref-1 inhibitors, this approach is feasible. Moreover, patient and healthcare system costs might be significantly lower with such a therapy, as office visits and injection procedures could be reduced.

In summary, we have explored Ref-1 inhibition as an ocular antiangiogenic therapy. We used two novel inhibitors of Ref-1 activity to demonstrate antiangiogenic effects *in vitro*, *ex vivo*, and in an animal model of CNV. The approach of targeting Ref-1 activity, and these small molecule inhibitors in particular, hold promise for development of therapies for treating diseases such as wet AMD, PDR, and ROP.

## Acknowledgements

We thank Rakshin Kharwadkar for assistance with proliferation assays.

## Authorship Contributions

*Participated in research design:* Pran Babu, Sishtla, Sulaiman, Park, Shetty, Shah, Fishel, Kelley, Corson

*Conducted experiments:* Pran Babu, Sishtla, Sulaiman, Park, Shetty, Shah

*Contributed new reagents or analytic tools:* Kelley, Wikel

*Performed data analysis:* Pran Babu, Sishtla, Sulaiman, Park, Shetty, Shah, Corson

*Wrote or contributed to the writing of the manuscript:* Pran Babu, Sishtla, Sulaiman, Park, Shetty, Shah, Fishel, Wikel, Kelley, Corson

## Footnotes

This work was supported by the Retina Research Foundation, the BrightFocus Foundation, the National Institutes of Health National Eye Institute [Grant R01EY025641], and an unrestricted grant from Research to Prevent Blindness, Inc. to T.W.C. Additional financial support for this work was provided by the National Cancer Institute [Grants R01CA138798 (to M.L.F. and M.R.K.), R01CA167291 (M.R.K.), R01CA205166 (M.R.K.)] and the National Institute of Neurological Disorders and Stroke [Grant R21NS091667 (to M.R.K.)], as well as financial support provided by the Earl and Betty Herr Professor in Pediatric Oncology Research, Jeff Gordon Children’s Foundation and the Riley Children’s Foundation (M.R.K.). APX3330, APX2009 and APX2014 were provided at no cost by Apexian Pharmaceuticals.

## Abbreviations

AMD: age-related macular degeneration
ANOVA: analysis of variance
AP-1: activator protein 1
APE1: apurinic/apyrimidinic endonuclease 1
APX2009: (*E*)-*N*,*N*-diethyl-2-((3-methoxy-1,4-dioxo-1,4-dihydronaphthalen-2-yl)methylene)pentanamide
APX2014: (*E*)-*N*-methoxy-2-((3-methoxy-1,4-dioxo-1,4-dihydronaphthalen-2-yl)methylene)pentanamide
APX3330: (2*E*)-2-[(4,5-dimethoxy-2-methyl-3,6-dioxo-1,4-cyclohexadien-1-yl)methylene]-undecanoic acid
ATCC: American Type Culture Collection
BSA: bovine serum albumin
CNV: choroidal neovascularization
DAPI: 4′,6-diamidine-2′-phenylindole
DMEM: Dulbecco’s modified Eagle’s medium
DMSO: dimethyl sulfoxide
EBM-2: endothelial basal medium 2
EdU: 5-ethynyl-2′-deoxyuridine
EGM-2: endothelial growth medium 2
EMSA: electrophoretic mobility shift assay
FDA: Food and Drug Administration
FA: fluorescein angiography
GCL: ganglion cell layer
GI_50_: median growth inhibitory concentration
GS-IB4: isolectin B4 from *Griffonia simplicifolia*
HIF-1α: hypoxia inducible factor 1α
HRECs: human retinal microvascular endothelial cells
INL: inner nuclear layer
L-CNV: laser-induced choroidal neovascularization
NF-κB: nuclear factor κ light-chain-enhancer of activated B cells
OCT: optical coherence tomography
ONL: outer nuclear layer
PBS: phosphate buffered saline
PDR: proliferative diabetic retinopathy
PKT: propylene glycol
Kolliphor HS15: Tween 80
Ref-1: reduction-oxidation factor 1
qRT-PCR: quantitative reverse transcription-polymerase chain reaction
ROP: retinopathy of prematurity
RP2D: recommended Phase II dose
RPE: retinal pigment epithelium
STAT3: signal transducer and activator of transcription 3
TUNEL: terminal deoxynucleotidyl transferase dUTP nick end labeling
VEGF: vascular endothelial growth factor
VLDLR: very low density lipoprotein receptor

## References

Basavarajappa HD, Lee B, Fei X, Lim D, Callaghan B, Mund JA, Case J, Rajashekhar G, Seo SY and Corson TW (2014) Synthesis and mechanistic studies of a novel homoisoflavanone inhibitor of endothelial cell growth. PLoS One 9:e95694.

Basavarajappa HD, Sulaiman RS, Qi X, Shetty T, Sheik Pran Babu S, Sishtla KL, Lee B, Quigley J, Alkhairy S, Briggs CM, Gupta K, Tang B, Shadmand M, Grant MB, Boulton ME, Seo SY and Corson TW (2017) Ferrochelatase is a therapeutic target for ocular neovascularization. EMBO Mol Med 9:786–801.

Biswas A, Khanna S, Roy S, Pan X, Sen CK and Gordillo GM (2015) Endothelial cell tumor growth is Ape/ref-1 dependent. Am J Physiol Cell Physiol 309:C296–307.

Campochiaro PA (2013) Ocular neovascularization. J Mol Med 91:311–321.

Cardoso AA, Jiang Y, Luo M, Reed AM, Shahda S, He Y, Maitra A, Kelley MR and Fishel ML (2012) APE1/Ref-1 regulates STAT3 transcriptional activity and APE1/Ref-1-STAT3 dual-targeting effectively inhibits pancreatic cancer cell survival. PLoS One 7:e47462

Chiarini LB, Freitas FG, Petrs-Silva H and Linden R (2000) Evidence that the bifunctional redox factor / AP endonuclease Ref-1 is an anti-apoptotic protein associated with differentiation in the developing retina. Cell Death Differ 7:272–281.

Day S, Acquah K, Mruthyunjaya P, Grossman DS, Lee PP and Sloan FA (2011) Ocular complications after anti-vascular endothelial growth factor therapy in Medicare patients with age-related macular degeneration. Am J Ophthalmol 152:266–272.

Evans AR, Limp-Foster M and Kelley MR (2000) Going APE over ref-1. Mutat Res 461:83–108.

Falavarjani KG and Nguyen QD (2013) Adverse events and complications associated with intravitreal injection of anti-VEGF agents: a review of literature. Eye 27:787–794.

Fishel ML, Colvin ES, Luo M, Kelley MR and Robertson KA (2010) Inhibition of the redox function of APE1/Ref-1 in myeloid leukemia cell lines results in a hypersensitive response to retinoic acid-induced differentiation and apoptosis. Exp Hematol 38:1178–1188.

Fishel ML, Jiang Y, Rajeshkumar NV, Scandura G, Sinn AL, He Y, Shen C, Jones DR, Pollok KE, Ivan M, Maitra A and Kelley MR (2011) Impact of APE1/Ref-1 redox inhibition on pancreatic tumor growth. Mol Cancer Ther 10:1698–1708.

Fishel ML, Wu X, Devlin CM, Logsdon DP, Jiang Y, Luo M, He Y, Yu Z, Tong Y, Lipking KP, Maitra A, Rajeshkumar NV, Scandura G, Kelley MR and Ivan M (2015) Apurinic/apyrimidinic endonuclease/redox factor-1 (APE1/Ref-1) redox function negatively regulates NRF2. J Biol Chem 290:3057–3068.

Forsythe JA, Jiang BH, Iyer NV, Agani F, Leung SW, Koos RD and Semenza GL (1996) Activation of vascular endothelial growth factor gene transcription by hypoxia-inducible factor 1. Mol Cell Biol 16:4604–4613.

Grossniklaus HE, Kang SJ and Berglin L (2010) Animal models of choroidal and retinal neovascularization. Prog Retinal Eye Res 29:500–519.

Jedinak A, Dudhgaonkar S, Kelley MR and Sliva D (2011) Apurinic/Apyrimidinic endonuclease 1 regulates inflammatory response in macrophages. Anticancer Res 31:379–385.

Jiang A, Gao H, Kelley MR and Qiao X (2011) Inhibition of APE1/Ref-1 redox activity with APX3330 blocks retinal angiogenesis in vitro and in vivo. Vision Res 51:93–100.

Kelley MR, Georgiadis MM and Fishel ML (2010) DNA Repair & Redox Signaling, in The Tumor Microenvironment (Teicher BA ed) pp 133–168, Springer Science+Business Media, Framingham.

Kelley MR, Georgiadis MM and Fishel ML (2012) APE1/Ref-1 role in redox signaling: translational applications of targeting the redox function of the DNA repair/redox protein APE1/Ref-1. Curr Mol Pharmacol 5:36–53.

Kelley MR, Luo M, Reed A, Su D, Delaplane S, Borch RF, Nyland RL, 2nd, Gross ML and Georgiadis MM (2011) Functional analysis of novel analogues of E3330 that block the redox signaling activity of the multifunctional AP endonuclease/redox signaling enzyme APE1/Ref-1. Antioxid Redox Signal 14:1387–1401.

Kelley MR and Wikel JH (2015) Quinone compounds for treating Ape1 mediated diseases. US Patent 9,193,700.

Kelley MR, Wikel JH, Guo C, Pollok KE, Bailey BJ, Wireman R, Fishel ML and Vasko MR (2016) identification and characterization of new chemical entities targeting apurinic/apyrimidinic endonuclease 1 for the prevention of chemotherapy-induced peripheral neuropathy. J Pharmacol Exp Ther 359:300–309.

Lando D, Pongratz I, Poellinger L and Whitelaw ML (2000) A redox mechanism controls differential DNA binding activities of hypoxia-inducible factor (HIF) 1alpha and the HIF-like factor. J Biol Chem 275:4618–4627.

Li L, Cheung SH, Evans EL and Shaw PE (2010) Modulation of gene expression and tumor cell growth by redox modification of STAT3. Cancer Res 70:8222–8232.

Li Y, Liu X, Zhou T, Kelley MR, Edwards P, Gao H and Qiao X (2014a) Inhibition of APE1/Ref-1 redox activity rescues human retinal pigment epithelial cells from oxidative stress and reduces choroidal neovascularization. Redox Biol 2:485–494.

Li Y, Liu X, Zhou T, Kelley MR, Edwards PA, Gao H and Qiao X (2014b) Suppression of choroidal neovascularization through inhibition of APE1/Ref-1 redox activity. Invest Ophthalmol Vis Sci 55:4461–4469.

Logsdon DP, Grimard M, Luo M, Shahda S, Jiang Y, Tong Y, Yu Z, Zyromski N, Schipani E, Carta F, Supuran CT, Korc M, Ivan M, Kelley MR and Fishel ML (2016) Regulation of HIF1alpha under hypoxia by APE1/Ref-1 impacts CA9 expression: Dual targeting in patient-derived 3D pancreatic cancer models. Mol Cancer Ther 15:2722–2732.

Lou D, Zhu L, Ding H, Dai HY and Zou GM (2014) Aberrant expression of redox protein Ape1 in colon cancer stem cells. Oncol Lett 7:1078–1082.

Luo M, Delaplane S, Jiang A, Reed A, He Y, Fishel M, Nyland RL, 2nd, Borch RF, Qiao X, Georgiadis MM and Kelley MR (2008) Role of the multifunctional DNA repair and redox signaling protein Ape1/Ref-1 in cancer and endothelial cells: small-molecule inhibition of the redox function of Ape1. Antioxid Redox Signal 10:1853–1867.

Luo M, Zhang J, He H, Su D, Chen Q, Gross ML, Kelley MR and Georgiadis MM (2012) Characterization of the redox activity and disulfide bond formation in apurinic/apyrimidinic endonuclease. Biochemistry 51:695–705.

Lux A, Llacer H, Heussen FM and Joussen AM (2007) Non-responders to bevacizumab (Avastin) therapy of choroidal neovascular lesions. Br J Ophthalmol 91:1318–1322.

McIlwain DW, Fishel ML, Boos A, Kelley MR and Jerde TJ (2018) APE1/Ref-1 redox-specific inhibition decreases survivin protein levels and induces cell cycle arrest in prostate cancer cells. Oncotarget 9:10962–10977.

McNeill DR and Wilson DM, 3rd (2007) A dominant-negative form of the major human abasic endonuclease enhances cellular sensitivity to laboratory and clinical DNA-damaging agents. Mol Cancer Res 5:61–70.

Nishi T, Shimizu N, Hiramoto M, Sato I, Yamaguchi Y, Hasegawa M, Aizawa S, Tanaka H, Kataoka K, Watanabe H and Handa H (2002) Spatial redox regulation of a critical cysteine residue of NF-kappa B in vivo. J Biol Chem 277:44548–44556.

Poor SH, Qiu Y, Fassbender ES, Shen S, Woolfenden A, Delpero A, Kim Y, Buchanan N, Gebuhr TC, Hanks SM, Meredith EL, Jaffee BD and Dryja TP (2014) Reliability of the mouse model of choroidal neovascularization induced by laser photocoagulation. Invest Ophthalmol Vis Sci 55:6525–6534.

Prasad PS, Schwartz SD and Hubschman JP (2010) Age-related macular degeneration: current and novel therapies. Maturitas 66:46–50.

Seo YR, Kelley MR and Smith ML (2002) Selenomethionine regulation of p53 by a ref1-dependent redox mechanism. Proc Natl Acad Sci U S A 99:14548–14553.

Shah F, Logsdon D, Messmann RA, Fehrenbacher JC, Fishel ML and Kelley MR (2017) Exploiting the Ref-1-APE1 node in cancer signaling and other diseases: from bench to clinic. NPJ Precis Oncol 1: 19.

Su DG, Delaplane S, Luo M, Rempel DL, Vu B, Kelley MR, Gross ML and Georgiadis MM (2011) Interactions of APE1 with a redox inhibitor: Evidence for an alternate conformation of the enzyme. Biochemistry 50:82–92.

Sulaiman RS, Merrigan S, Quigley J, Qi X, Lee B, Boulton ME, Kennedy B, Seo SY and Corson TW (2016) A novel small molecule ameliorates ocular neovascularisation and synergises with anti-VEGF therapy. Sci Rep 6:25509.

Sulaiman RS, Quigley J, Qi X, O’Hare MN, Grant MB, Boulton ME and Corson TW (2015) A simple optical coherence tomography quantification method for choroidal neovascularization. J Ocul Pharmacol Ther 31:447–454.

Vasko MR, Guo C, Thompson EL and Kelley MR (2011) The repair function of the multifunctional DNA repair/redox protein APE1 is neuroprotective after ionizing radiation. DNA Repair (Amst) 10:942–952.

Wenzel AA, O’Hare MN, Shadmand M and Corson TW (2015) Optical coherence tomography enables imaging of tumor initiation in the TAg-RB mouse model of retinoblastoma. Mol Vis 21:515–522.

Xanthoudakis S and Curran T (1992) Identification and characterization of Ref-1, a nuclear protein that facilitates AP-1 DNA-binding activity. EMBO J 11:653–665.

Xanthoudakis S, Miao G, Wang F, Pan YC and Curran T (1992) Redox activation of Fos-Jun DNA binding activity is mediated by a DNA repair enzyme. EMBO J 11:3323–3335.

Yan T, Venkat P, Chopp M, Zacharek A, Yu P, Ning R, Qiao X, Kelley MR and Chen J (2018) APX3330 promotes neurorestorative effects after stroke in type one diabetic rats. Aging Dis, 10.14336/AD.2017.1130.

Zhang J, Luo M, Marasco D, Logsdon D, LaFavers KA, Chen Q, Reed A, Kelley MR, Gross ML and Georgiadis MM (2013) Inhibition of apurinic/apyrimidinic endonuclease I’s redox activity revisited. Biochemistry 52:2955–2966.

Zou GM, Karikari C, Kabe Y, Handa H, Anders RA and Maitra A (2009) The Ape-1/Ref-1 redox antagonist E3330 inhibits the growth of tumor endothelium and endothelial progenitor cells: therapeutic implications in tumor angiogenesis. J Cell Physiol 219:209–218.

